# Neuropixels 1536 Channel Quad Base probe reveals brain-wide communication underlying flexible sensorimotor sequences

**DOI:** 10.64898/2026.07.23.740388

**Authors:** Yi-Ting Chang, Rajan Dasgupta, Mingyuan Dong, Jennifer Colonell, Bill Karsh, Alexandra Cheng, Ruben Hermans, Hasan Mahmud-Ul, Carolina Mora Lopez, Jan Putzeys, Marleen Welkenhuysen, Barundeb Dutta, Austin R. Graves, Daniel O’Connor, Timothy Harris

## Abstract

The brain orchestrates billions of neurons spanning hundreds of regions to drive behavior. High-density electrophysiology has provided insights into this process, but monitoring brain-wide patterns of spiking activity demands new tools. Here we present Neuropixels 2.0 Quad Base, a 1536-channel probe with four times the simultaneous recording capacity of state-of-art Neuropixels 2.0. Using two Quad Base probes, we performed 3072-channel recordings in mice trained to produce complex sequences of goal-directed licks. We decoded behavioral kinematics from neural populations across more than 20 brain regions and identified the spatiotemporal emergence of motor decision signals. The probes’ high channel count further revealed expanded inter-neuronal functional subnetworks and latent dynamics across multiple areas—features significantly undersampled when channels were subsampled to mimic standard Neuropixels 2.0. Neuropixels 2.0 Quad Base offers a powerful tool to explore neural encoding of cognition and behavior at grand scale.

## Introduction

A central goal of neuroscience is to understand how behavior is driven by neural activity across scales: from spiking of single neurons to plasticity within neural circuits and ultimately whole-brain systems.^1,2^ Over the past half century, many techniques have been used to observe neural activity in behaving rodents. In vivo electrophysiology (e.g. tetrodes, Utah arrays, and silicon probes) is widely used to record from dozens to hundreds of single neurons either at the probe tip or in tight columns of brain tissue along the probe shank.^3–5^ In vivo two-photon imaging offers high spatial resolution and expansive coverage of the dorsal cortex but lacks temporal resolution to reveal neural firing and is not optimal for broad coverage of deeper structures.^6–8^ fMRI can achieve high spatial coverage and reveal synchronous changes in activity across the brain, but it lacks the spatial and temporal resolution to reveal cellular mechanisms of neural computation.^9^ Despite a wealth of data demonstrating how behavior relates to neural activity within individual regions, our knowledge of how these processes work in concert across the brain is extremely limited. Elucidating these global mechanisms of cognition demands recording techniques capable of achieving high spatial and temporal resolution synchronously from across the brain.

Since their inception a decade ago, Neuropixels probes have revolutionized systems neuroscience by significantly enhancing the scale of neural recordings. While tetrode arrays and silicon probes offer between 4-64 recording sites, Neuropixels 1.0^4^ and 2.0^4,10^ probes (NP 1.0 and NP 2.0) offer 384 channels without compromising the spatial and temporal resolution required to resolve single-neuron spiking. These probes have been used by thousands of labs to record neural activity in behaving rodents, primates, and even humans. To optimize the accuracy of spike sorting, most Neuropixels experiments confine the 384 user-defined recording channels to either a single brain region or at most a few regions, each surveyed with at least 100 recording sites. While brain-wide recordings of activity have been made using sequential insertions of several Neuropixels 2.0 probes,^11,12^ there remains an urgent need for probes with higher channel counts capable of dense simultaneous recordings across brain areas.

Here, we present Neuropixels 2.0 Quad Base (NP QB): a new electrophysiology probe with 1536 simultaneous recording channels, a 4-fold increase in recording capacity compared to current state-of-the-art NP 2.0 probes. Neuropixels Quad Base probes are the first tool that enables high-density recording along the full depth of the probe, avoiding the tradeoff between spatial coverage and recording density that limits current generations of Neuropixels probes. Significantly, the increased channel count of Quad Base probes unlocks fundamentally new experimental avenues to explore brain-wide encoding of behavior. To demonstrate this utility with Neuropixels Quad Base probes, we recorded dense neural activity simultaneously across a distributed set of brain regions in mice performing a flexible, sensory-driven sequence licking task. These data revealed novel brain-wide ensembles involved in sensory-motor transformations and elucidated with unprecedented spatiotemporal detail the emergence of motor reprogramming signals. Further, these brain-wide data enable fundamentally new analyses to identify functional subnetworks and inter-regional latent dynamics. Ultimately, the brain-wide recordings enabled by the new Neuropixels Quad Base probes represent the first step along a path to exploring how encoding of complex behaviors is distributed across the brain.

## Results

### Neuropixels Quad Base quadruples the recording capacity of standard Neuropixels

Building upon the standard Neuropixels 2.0 (NP 2.0) platform,^10^ which provides 384 recording channels for high-density electrophysiology, we introduce a significant advancement: the Neuropixels Quad Base (NP QB). This design quadruples the recording capacity to an unprecedented 1,536 channels (Fig. 1a), maintaining noise and gain characteristics comparable to those of the NP 2.0 (Fig. 1b and Fig. S1a; see Ye, et al.^13^ for details of the measurements; NP 2.0 noise specification is 6.8 μV_rms_). While retaining the four-shank geometry (10 mm length) and 5,120 low-impedance TiN sites of the NP 2.0, the NP QB incorporates a scaled probe base (width: 10.2 mm in NP QB, 3.5 mm in NP 2.0) and headstage (NP QB: 14 x 18 mm, NP 2.0: 10 x 14 mm) to accommodate the expanded electronics required for this fourfold increase in capacity. This architecture enables simultaneous, large-scale recordings across broadly distributed neuronal populations within a single experimental preparation (Fig. 1c), providing a unique window to investigate complex circuits with exceptional spatial and temporal resolution.

**Figure 1:**
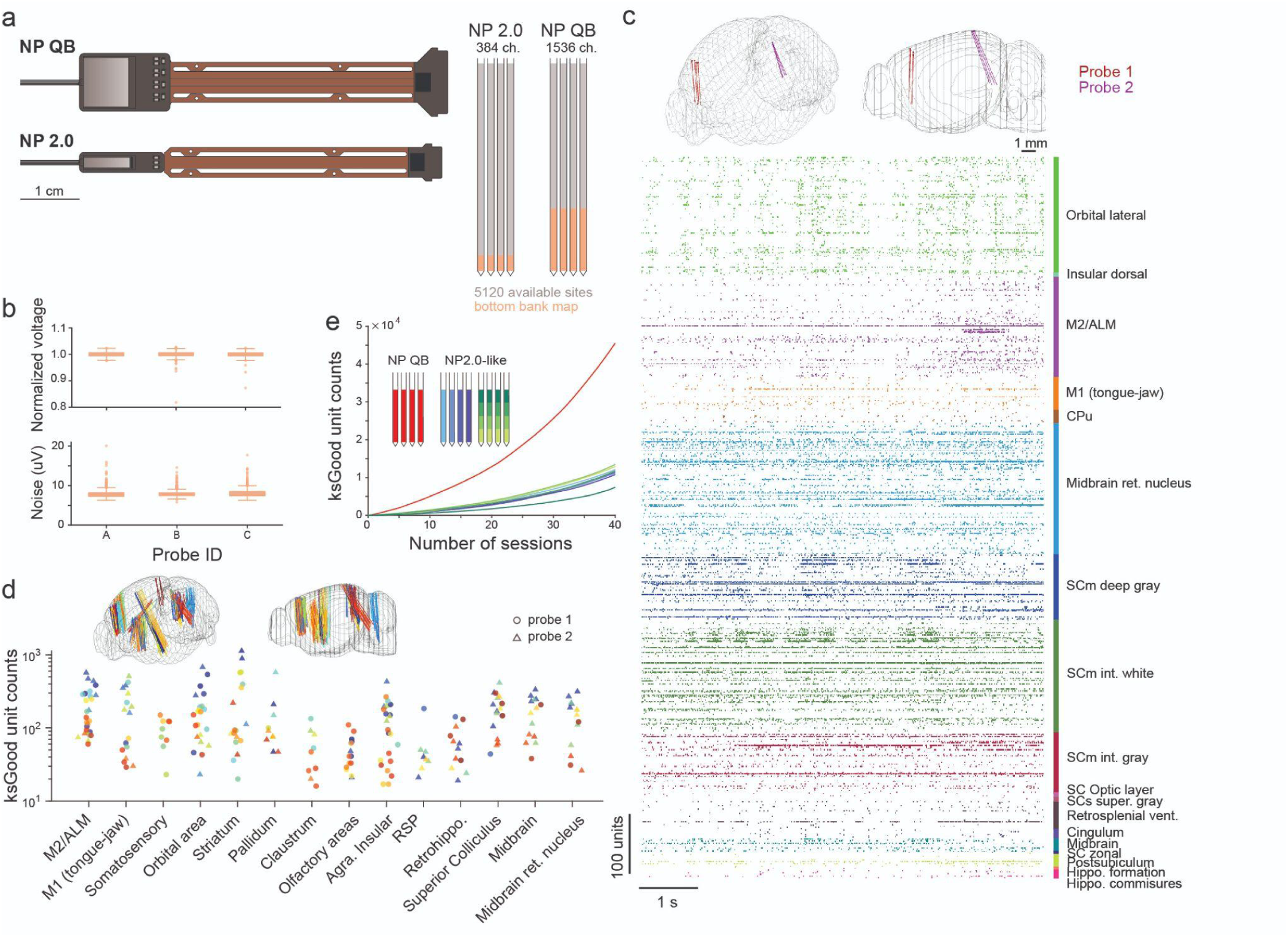
Neuropixels 2.0 Quad Base quadruples the recording capacity of standard probes. **a**, Comparison of the Neuropixels 2.0 (NP 2.0) and NP Quad Base (NP QB) probe architectures. The NP QB maintains the standard shank layout but increases the channel count from 384 to 1536 for large-scale simultaneous recording. **b**, *In vitro* measurements of noise (integrated from 300-10000Hz, lower panel) and gain uniformity (upper panel) for three NP QB probes. These values are comparable to the NP2.0 noise specification of 6.8 μV_rms_. **c**, Example spike rasters recorded simultaneously using two NP QB probes in an awake, head-fixed mouse. See brain region abbreviations in Table S1. **d**, Quantification of neuron yields across 14 selected brain regions (n = 40 recordings from 6 mice). Each color represents a single recording session with two NP QB probes (probe 1: circle; probe 2: triangle). **e**, Comparison of neuron yield between NP QB (red) and data resampled to simulate standard NP 2.0 channel maps (NP2.0-like, blue and green).

To validate the NP QB platform, we performed dual-probe recordings in awake, head-fixed mice (Fig. 1c–e and S1). We obtained activity from large neural populations across 40 sessions in 6 mice (Fig. 1d and Table S1), achieving an average yield of over a thousand Kilosort good^14^ neurons per session (n = 1139 ± 94 units). To benchmark the NP QB against the standard configuration, we generated an ’NP2.0-like’ control dataset by computationally resampling the NP QB data to mimic the channel map and capacity of a standard NP 2.0 probe (n = 285 ± 9 units, Methods). Thanks to its four-fold channel capacity, the NP QB dataset achieves equivalent neural population sizes in significantly fewer sessions than the NP2.0-like dataset (Fig. 1e). These results demonstrate that NP QB probes substantially increase the number of simultaneously isolated units, providing a critical tool for scaling research toward whole-circuit dynamics.

### NP QB probes enable brain-wide decoding of task-related variables in a skilled licking task

Our next undertaking was to validate the quality of the data generated by NP QB. We turned to a previously published skilled “sequence licking” task in mice, boasting a rich repertoire of associated neural factors^15^ (Fig. 2a-b). Thirsty mice carried out a series of directed licks following an auditory go cue, going through 7 port locations from one side to the other, and received a water reward at the end (Fig. 2a). After a random interval, the next trial went through the same port locations but in reverse. While mice carried out the task, we captured two views with high-speed video (bottom and side, Fig. 2b). From the video frames, we derived instantaneous measurements of a number of task and behavioral variables, including the locations of tongue and jaw tips in 3D space, the length of the tongue and its azimuthal angle from midline (Fig. 2b-c; see Methods for a full accounting of all variables).

**Figure 2:**
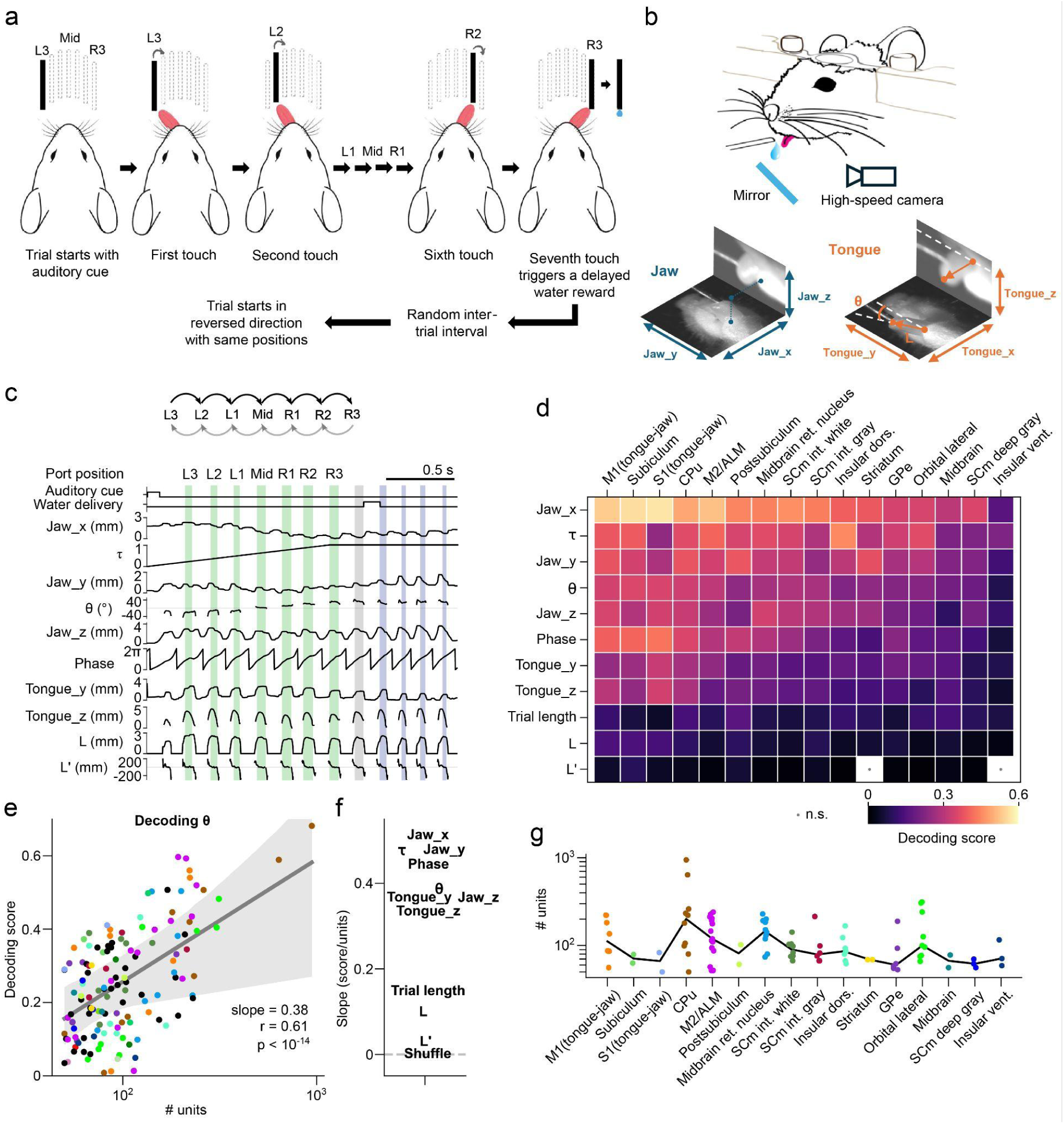
Sequential licking task and decoding kinematic variables from dense brain-wide recordings. **a**, Schematic of the sequential licking task. Following a 100-ms auditory go cue, mice guided their tongues to contact a motorized lick port across a stereotyped sequence of positions (L3–R3). Completion of the sequence triggered delayed water delivery. After a random inter-trial interval, the next trial went through the same port locations but in the opposite direction. **b**, High-speed videography configuration and kinematic coordinate systems. Simultaneous bottom- and side-view recordings (via mirror) were used to track jaw position and tongue posture in three dimensions. Jaw position is represented by Cartesian coordinates (Jaw_x, Jaw_y, Jaw_z). Tongue posture was parameterized by the azimuthal angle in the horizontal plane (θ), and tongue length (L). **c**, Example trial showing task events and behavioral variables aligned to the ongoing lick sequence. Kinematic variables were extracted from high-speed video; phase was computed from Jaw_z, and τ denotes relative sequence time. Shaded areas indicate periods of port contact: green for ‘drive’ licks that are part of the required sequence, gray for a lick prior to water consumption, and blue for consumptive licks. Port positions at contact during sequence execution are indicated above. **d**, Heatmap showing mean population decoding performance for behavioral and task parameters from population activity in selected brain regions (see Methods). Both behavioral/task parameters and brain regions are ordered according to their average decoding scores across all regions and parameters, respectively. As indicated, decoding performance was not significantly different from shuffled controls in some cases. **e**, Scatterplot showing θ-decoding score versus number of single units recorded in single sessions from different regions. Only recordings with 50 or more isolated single units were included. Colored dots indicate recordings from different brain regions (same color scheme as in (**g**)). Black dots indicate other brain regions not in (**g**). Solid grey line shows mean linear fit to the data, error shading represents 95% hierarchical bootstrap confidence intervals. Slope and Pearson correlation statistics are shown. **f**, Slopes of linear fits to decoding score versus unit number scatterplots, for different decoded variables. **g**, Number of units in each recording for the brain regions in (d). Black line shows region medians.

To validate the quality of isolated spikes, we began by linearly decoding the concurrently observed behavioral and task variables from the neural data obtained from each brain region. We restricted these analyses to recordings that included a minimum of fifty well-isolated single units from each region across a minimum of two subjects. All the variables could be decoded significantly above chance levels from most brain regions, showing that it was possible to extract useful latent structures from the data acquired by NP QB probes from a wide variety of regions (Fig. 2d). However, the degree of decodability varied between variables and regions - sensorimotor cortical regions in particular showed especially good decoding performance.

As decoding models generally improve when there is more information in the data for the model to exploit,^16^ decoding performance should improve with the inclusion of more units.^17^ Indeed, the ability to decode was strongly correlated with the number of units that were captured in each recording (Fig. 2e-f), demonstrating the importance of the kind of high-density recordings that are enabled by NP QB. The trend of better decoding performance for more units held for all variables considered, but was stronger for variables with overall better decoding (Fig. 2f). Interestingly, greater overall decodability from certain regions could not be readily explained by discrepancies in unit counts (Fig. 2g). Together, these analyses showed that NP QB recordings produced high-quality data from which meaningful biological signals could be reliably extracted.

### Emergence of critical motor decision points across brain regions

To encourage flexible motor execution, in addition to “standard” sequence trials in which mice licked straight through the 7 port locations, each session included a random subset of interleaved “backtracking” sequence trials (30% probability). In these trials, the port moved backwards after the lick at Mid position (Fig. 3a, top and middle). The unexpected backward movement of the port caused the mice to make a “missed lick” in the lick cycle following lick port contact at the Mid position. Expert mice learned to rapidly branch their ongoing motor execution to recover the port (at L2 or R2) and finish the sequence (Fig. 3a, bottom). As this required a drastic sensory-driven reorientation of ongoing motor programs, state space trajectories of population activity showed rapid and dramatic divergence between standard and backtracking sequences following the lick at Mid (Fig. 3b).

**Figure 3:**
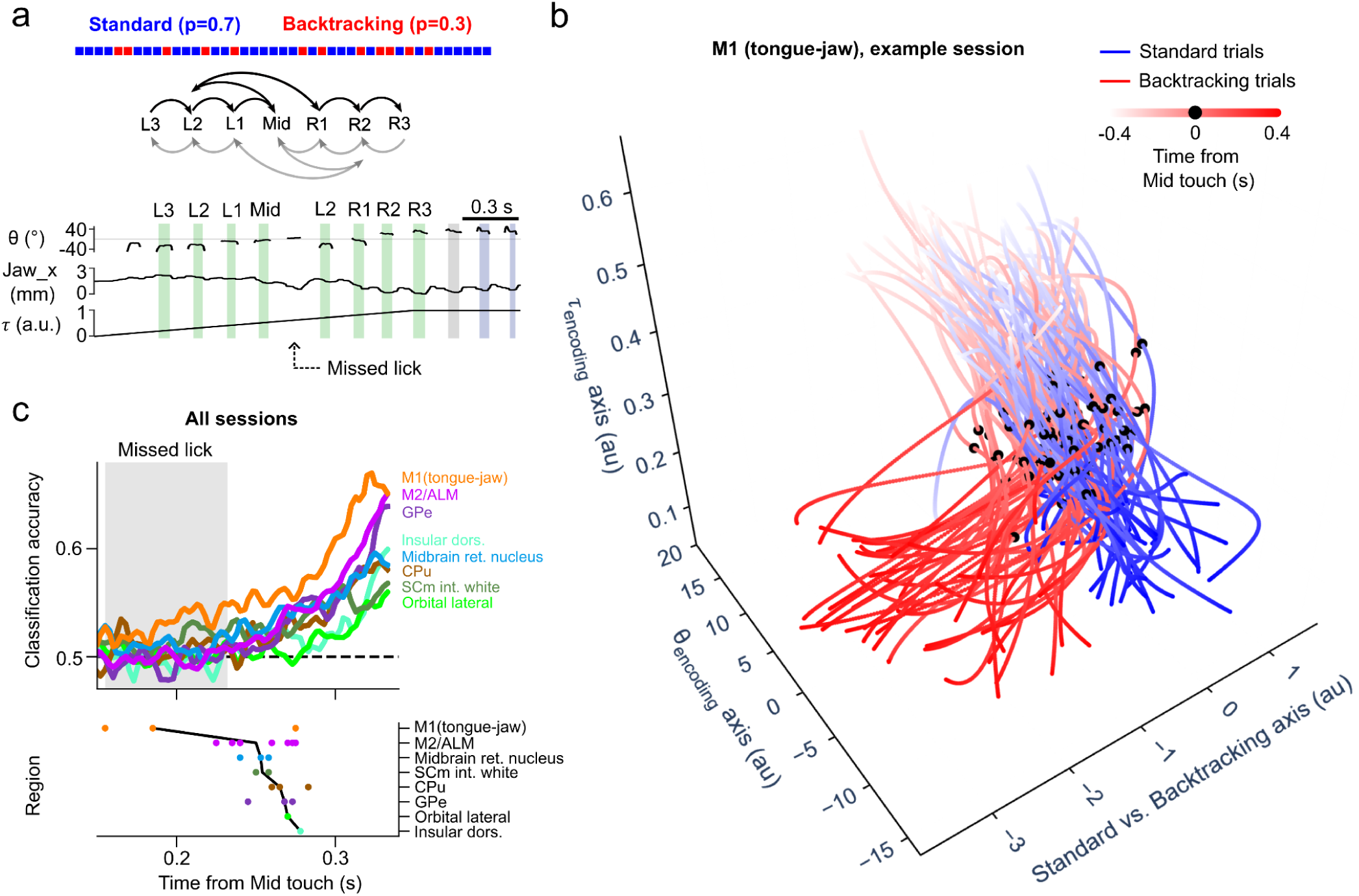
Detecting the first occurrence of motor branching decisions from simultaneous brain-wide recordings. **a**, Top, schematic of task structure, showing randomly interspersed trials with backtracking sequences. Middle, schematic showing lick port movements in backtracking sequences in each direction. Bottom, tongue angle θ and Jaw_x traces from an example backtracking trial. **b**, State space trajectories of M1 (tongue-jaw) population activity from individual standard (blue) and backtracking (red) trials in a single session. Activity was projected onto the latent dimensions that best predicted τ and θ (“τ-encoding” and “θ-encoding” axes, see Methods) and the dimension that best discriminated between the population activity in standard and backtracking trials after the lick at Mid. **c**, Top, time-series of median classification performance of classifiers trained to distinguish between standard and backtracking trials using the population activity in each time-bin. Median classification performance values on a shuffled dataset were subtracted. The period of the missed lick is indicated with gray shading. Bottom, time of start of significant trial-type classification performance for individual sessions and regions. Black line shows the region medians.

Recent studies have identified subcortical areas like the superior colliculus as being crucial for rapid touch-guided adjustments to lick angles.^18^ As multiple behavioral and task variables were represented widely across brain areas in our task (Fig. 2d), we wondered where the decision to commit to backtracking first emerged following the detection of the sensory hallmarks of backtracking trials. We took advantage of the simultaneity of the recordings from a wide range of brain areas enabled by the NP QB probes to answer this question. We trained linear classifiers to distinguish between standard and backtracking trials using the population activity in each brain region during each 2.5 ms bin (Methods). Among the brain regions queried, classification performance began to improve from chance levels first in M1 (tongue-jaw subregion), followed by the anterolateral motor cortex, ALM (Fig. 3c). By contrast, classification performance lagged in the collicular regions, suggesting that in our task (where motor corrections were affected by the absence of a predicted touch input rather than touch itself), cortical regions were the principal drivers of flexible motor control. Thus, dense brain-wide recordings using NP QB probes enabled interrogations of the time courses of rapid shifts in brain states during active behaviors.

### NP QB probes enable the discovery of brain-wide functional sub-networks

Next, we asked whether NP QB probes could improve our ability to identify brain-wide functional subnetworks. We focused on two time periods within each trial in which the mouse was in distinct behavioral states. From each trial, we isolated the one second long “Licking” period centered on the time of port contact in the Mid position, and the “Not licking” period that took place 2-3 seconds after the delivery of water at the end of the trial (Fig. 4a). Then, we looked for significant Granger causation between each pair of single units obtained from different brain regions in the recording (Methods). The total number of Granger causal links was much higher when the mouse was licking (Fig. 4b), consistent with widespread sensory and motor signals associated with sequence execution. The widespread reductions in the numbers of Granger causal links during the not licking period could not be readily explained by differences in firing rates in the two states (Fig. S2).

**Figure 4:**
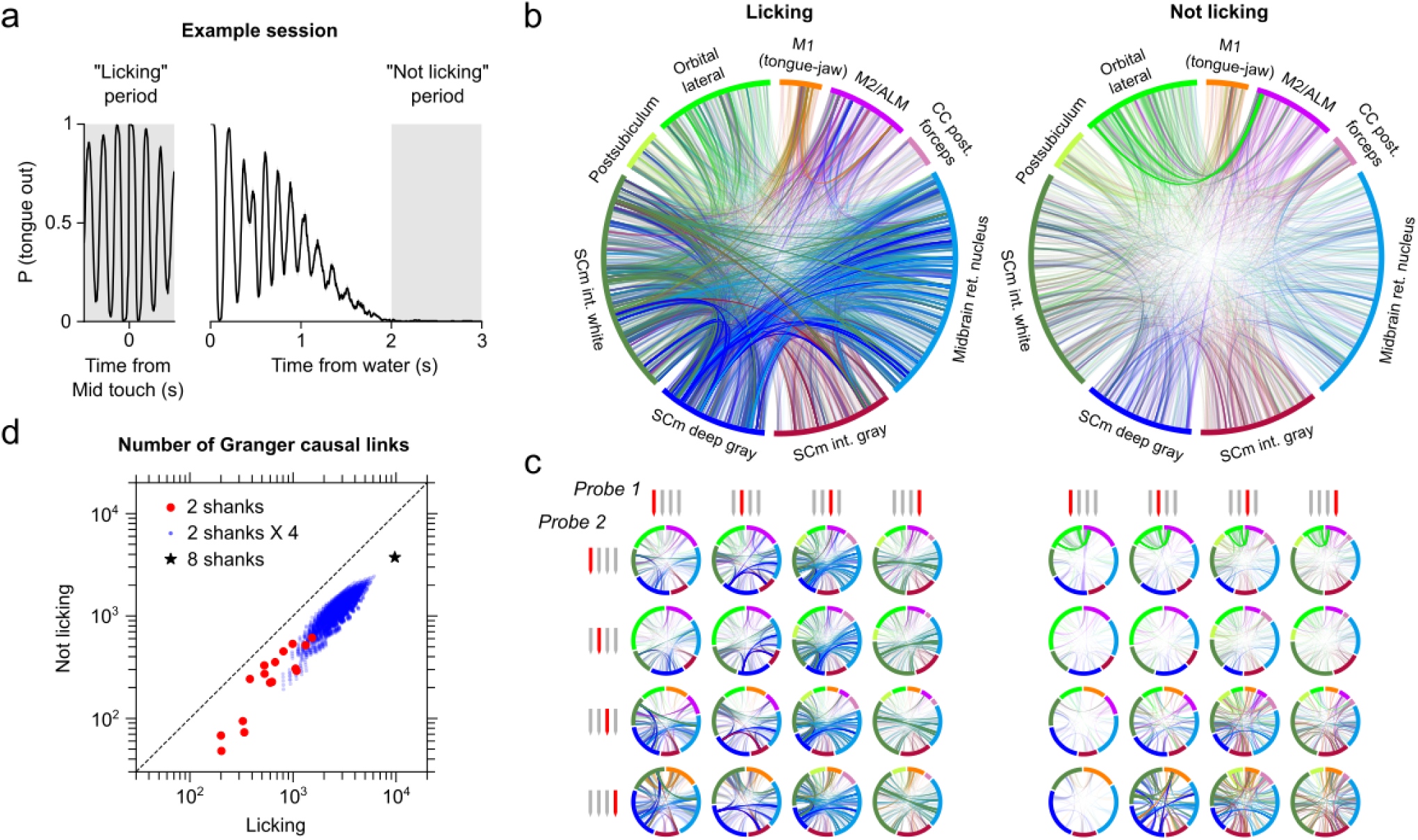
Neuropixels QB enables the detection of more functional subnetworks across brain areas. **a**, Probability of the tongue being out of the mouth during the 1-second long Licking and Not licking periods (indicated by grey shading) of all trials in an example session. **b**, Chord diagrams showing Granger causal links between individual units across brain regions during the two periods. The colors of individual links correspond to the region that yielded stronger causality as the Granger-cause, i.e. predictor, in each pair (see Methods). Strengths of links are indicated by their opacities. Links between units within the same region are hidden. **c**, Array of chord diagrams showing detected Granger causal links between units recorded in each of sixteen possible 2-shank combinations, representing mock 2-probe single-shank NP2.0-like experiments. Shanks being considered in each case are indicated in red. **d**, Scatterplot showing numbers of detected Granger-causal links. The number of links detected in a single 8-shank (dual NP QB, black star) recording exceeded the theoretical number of detected links from four consecutive 2-shank (dual NP2.0-like) recordings (2 shanks X 4, blue). Link numbers from individual 2-shank (dual NP2.0-like) recordings from (c) are indicated in red.

We explored how many of those links would have been identified in hypothetical dual single-shank NP2.0-like control experiments. We generated the control dataset by resampling single shanks from each of the two NP QB probes, and considered the number of Granger causal links formed between pairs of units recorded on each of the 4 X 4 possible pairs of shanks. There were dramatic qualitative differences between the subnetworks identified when only considering the units picked up on sub-sampled pairs of shanks (Fig. 4c), showing that the equivalent NP2.0-like experiment would have failed to identify a significant portion of brain-wide neural dynamics. We wondered if more experiments using NP2.0-like configurations would have sufficed to capture all the Granger causal links. Notably, the number of links identified by one dual NP QB probe recording (8 shanks) exceeded the number of links obtained from four consecutive NP2.0-like experiments (2 shanks X 4) despite sampling from the same total channel count, (Fig. 4d), since the consecutive NP2.0-like recordings would miss links between units picked up by different shanks of the same probe. Thus, NP QB facilitated the identification of many more functional subnetworks than would have been possible with the current state-of-the-art.

### NP QB probes uncover richer shared latent dynamics across distributed networks

The high channel count of the NP QB enables simultaneous recording of thousands of neurons across multiple regions—a capability critical for dissecting distributed brain networks. To investigate the large-scale dynamics occurring across these broad networks, we employed mDLAG^19,20^ (delayed latent across multiple groups) analysis to identify concurrent shared latent dimensions of activity (Fig. 5a).

**Figure 5:**
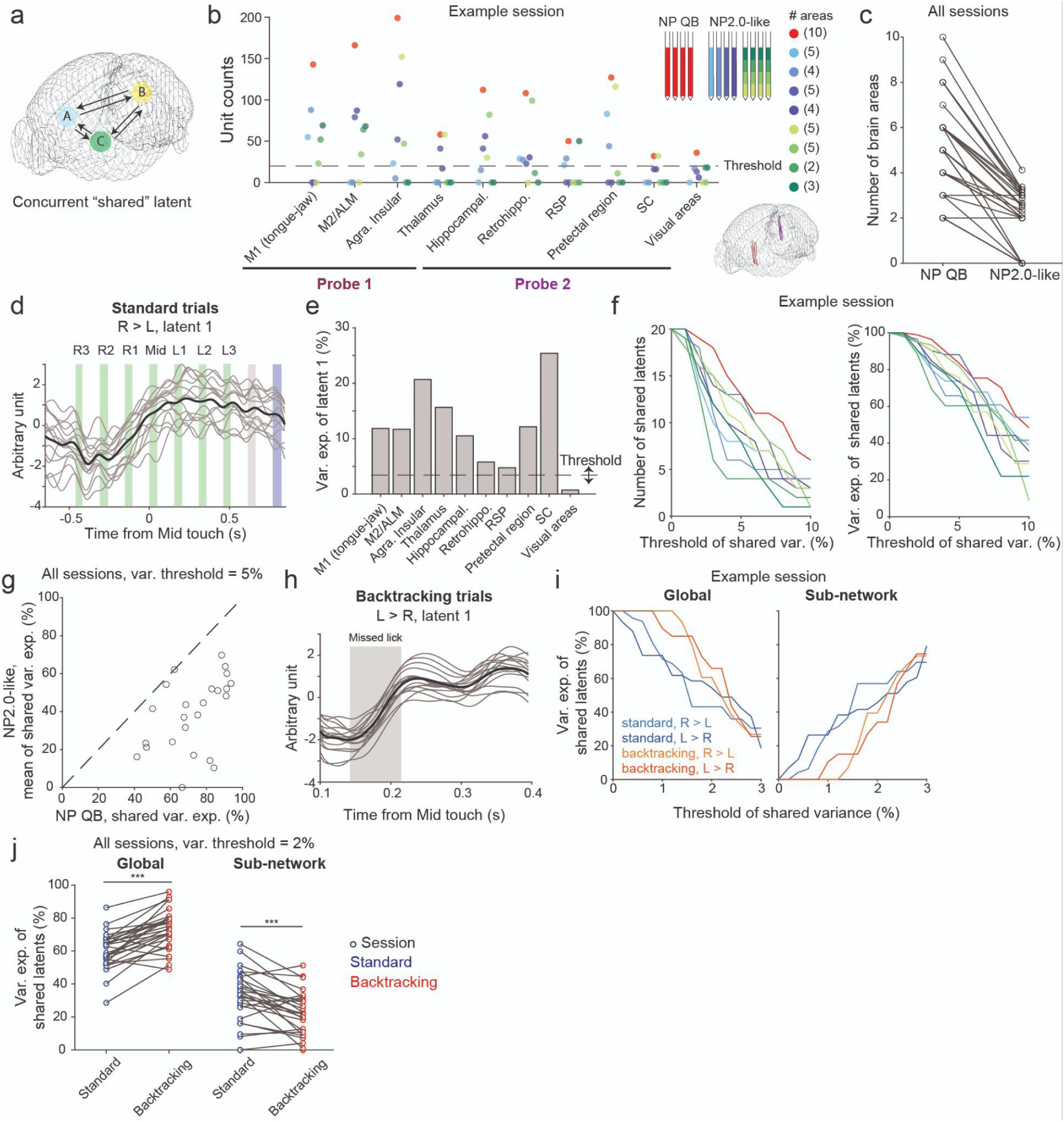
Neuropixels QB allows identification of richer shared latent dimensions across multiple brain areas. **a**, Schematic of mDLAG (delayed latent across multiple groups) illustrating the identification of shared latent dimensions across multiple brain areas. **b**, Comparison of the total number of units recorded in different brain areas using the NP QB probe (red) versus resampled NP2.0-like probes (blue and green) in a dual-probe recording session. A minimum of 20 units was required for a brain area to be included in the mDLAG analysis (NP QB: 10 brain areas; NP2.0-like: 2-5 brain areas). **c**, Number of brain areas of NP QB and NP2.0-like probes in the mDLAG analysis. For each NP QB recording, the brain area counts for all corresponding NP2.0-like simulations are presented as an average. **d**, Example time series of latent dimensions identified from the NP QB recording (in **b**). The traces show 15 randomly selected trials (gray) and the mean across all trials (black) in the primary motor area (M1). Shaded areas indicate periods of port contact: green for drive licks, gray for a lick prior to water consumption, and blue for consumptive licks. **e**, Variance explained by an example latent dimension (in **d**) across different brain areas. A threshold was applied to determine if a latent dimension significantly contributed to the variance explained in a given brain area. **f**, Number of shared latents and variance explained by all shared latents across multiple significance thresholds. In an example session (**b**), the NP QB probe (red) identified more shared latents (left panel) and captured more variance (right panel) compared to NP2.0-like probes (blue and green). **g**, Shared variance explained of NP QB and NP2.0-like probes. For each session, NP QB’s variance explained is plotted against the mean of variance explained of the corresponding NP2.0-like simulations (variance threshold = 5%). N = 31 sessions. **h**, Example latent dimensions identified during sensory-guided motor program reorganization (backtracking trials). A shaded box indicates a missed lick. **i**, Variance explained by global and sub-network shared latent dimensions across multiple significance thresholds. Global latents are defined as those significantly contributing to the variance in all recorded brain areas; sub-network latents are significant in a subset of brain areas (more than 2 but fewer than the total number of areas). **j**, Global and sub-network shared variance in standard and backtracking trials. The global latents explain more variance in backtracking trials (red), whereas sub-network latents explain more variance in standard trials (blue).

Reliable mDLAG analysis requires an adequate sample size for statistical power, which we defined as a minimum of 20 single units per brain area. The NP QB increased the number of areas meeting this criterion compared to standard configurations (Fig. 5b-c, mean number of areas: 5 in NP QB, 2 in NP2.0-like), thereby providing the data volume necessary for robust, large-scale network analysis. Although mDLAG is an unsupervised method, the latents it uncovered exhibited functionally relevant temporal structure. For instance, Latent 1 (ranked by mean variance explained across all brain regions) displayed left-right selectivity corresponding to the overall direction of licking sequence progression (left-to-right or right-to-left sequences) and rhythmic modulation corresponding to lick cycles (Fig. 5d). The latents explained varying fractions of variance across recorded regions (Fig. 5e).

We next assessed whether the NP QB could capture a richer network structure than standard probes. To determine whether a specific latent dimension significantly contributed to the variance explained within a given brain area, we applied a significance threshold (Fig. 5e, dashed line), defining “shared latents” as those contributing to more than two brain areas.As expected, increasing this significance threshold led to a decline in the number of identified shared latents (Fig. 5f, left). However, in an example session, the number of shared latents identified via NP QB was consistently higher across a range of significance thresholds compared to NP2.0-like simulations (Fig. 5f, left). This indicates that expanding unit yield per region provides the population sizes necessary to robustly track shared latent subspaces across distributed brain regions. Moreover, these shared latents in the NP QB accounted for a higher proportion of overall variance (Fig. 5f, right), suggesting they capture more meaningful shared population dynamics. Across all recording sessions using a conservative threshold of 5% variance explained, the majority of NP QB recordings demonstrated higher network variance explained than the mean of their corresponding NP2.0-like simulations (Fig. 5g, see Fig. S3 for consistent results across alternative thresholds). Overall, these results indicate that the NP QB captures complex inter-regional covariance structures and organizational principles that remain obscured by lower-yield recording technologies.

### The brain network shifts to a global processing state during motor program reorganization

Finally, we examined network reorganization during the sequence licking task. We investigated whether the sensory-driven switch of motor programs in backtracking trials recruited specific latent dynamics (Fig. 5h). While the total number of shared dimensions remained stable between standard and backtracking trials (Fig. S4a, number of shared dimensions: 15.3 ± 0.1 in standard, 16.1 ± 0.2 in backtracking, pair-t test p = 0.43), the composition of these dimensions shifted significantly (Fig. 5i–j and S4).

We classified latents as either Global (contributing significantly to variance in all recorded areas) or Sub-network (significant in >2 areas but fewer than the total). During backtracking, we observed a distinct transition in network organization: Global latents explained a significantly higher variance during backtracking compared to standard trials (Fig. 5j left, global variances: 60 ± 4% in standard, 73 ± 4% in backtracking, paired t-test p < 0.001, n = 29 sessions), whereas Sub-network latents explained less variance in backtracking trials (Fig. 5j right, sub-network variances: 31 ± 6% in standard, 22 ± 5% in backtracking, pair-t test p < 0.001, n = 29 sessions). These results suggest that sensory-driven motor switching necessitates a functional shift from more specialized local processing toward a highly integrated, whole-brain processing state.

## Discussion

We introduce the Neuropixels Quad Base (NP QB), a 1536-channel probe providing four-fold the channel capacity of Neuropixels 2.0. By deploying dual NP QB probes in mice, we simultaneously recorded over a thousand neurons, significantly expanding both unit yield and volumetric brain coverage (Fig. 1). Our results demonstrate that this increased channel count directly improves population decoding performance (Fig. 2) and enables the precise identification of the spatiotemporal emergence of motor decision signals (Fig. 3). Compared to simulated “NP 2.0-like” datasets, the NP QB facilitated the discovery of directed functional subnetworks via pairwise Granger causality analysis (Fig. 4) and identified inter-regional shared latents that account for a substantially greater proportion of neural variance (Fig. 5). Together, these findings establish the NP QB as a transformative tool for large-scale, brain-wide research.

The scaling of decoding accuracy with neuronal yield was consistent across all task variables,^17^ yet the improvement was most pronounced for variables with higher baseline decodability (Fig. 2e–f). This suggests that our targeted regions may preferentially encode these specific variables, or that the relevant information is more broadly distributed across the sampled populations. Furthermore, the sensitivity of these variables to population size likely reflects a higher underlying dimensionality, necessitating larger neuronal ensembles for accurate representation. In contrast, the plateauing or lower decoding performance observed for other variables may reflect that the signals are more redundant or constrained by the information-limiting correlations.^21–23^ It may also stem from non-linear encoding properties that are not fully captured by our linear decoding approaches.^24^ Future investigations involving recordings beyond the current target areas, as well as formal dimensionality^25,26^ and non-linear analyses^27^ will be essential to further elucidate the large-scale organization of these signals.

The large simultaneous recordings provided by the NP QB facilitate an investigation of neural communication across scales, ranging from fine-grained neuron-to-neuron interactions to large-scale coordination across multiple brain regions. We found that the NP QB identifies significantly more Granger causal links between neurons than simulated NP 2.0-like probes (Fig. 4), suggesting that high channel counts are essential for resolving directed functional connectivity that is otherwise lost to undersampling. Moreover, the NP QB reveals a richer landscape of inter-areal communication by capturing shared latent signals that explain a larger portion of neural variance (Fig. 5f–g), providing a more robust estimate of circuit-wide information flow. These results suggest that signals previously obscured by sparse sampling can be effectively recovered by scaling recording technology toward whole-circuit density.

A defining feature of our behavioral task is the requirement for mice to rapidly adapt their motor programs based on tactile feedback. We observed a temporal hierarchy in the emergence of motor decision signals,originating first in the primary motor cortex (M1 tongue/jaw), followed by the secondary motor cortex (M2/ALM), and finally the superior colliculus (SC) (Fig. 3c). This sequence suggests that motor cortical areas, rather than the colliculus, serve as the primary orchestrators of behavioral flexibility.^28,29^ While some studies have identified the colliculus as a principal driver of adaptive motor responses,^18^ this divergence may stem from task-specific demands. In our paradigm, motor corrections were triggered by the omission of a predicted touch input,^15^ whereas other studies often focus on corrections elicited by the presence of external tactile stimuli.^18^

Beyond the localized emergence of these signals, we observed a fundamental shift in brain network properties during motor reprogramming (Fig. 5i–j). The network transitioned from a state of specialized, local processing toward a highly integrated, brain-wide state. This global shift suggests that error signals—or the resulting motor correction commands—are propagated widely to coordinate the multiple regions required to modify an ongoing motor sequence. By capturing this transition across a massive population of neurons, the NP QB provides a unique window into how the brain balances local computation with global coordination to maintain behavioral flexibility.

**Figure S1:**
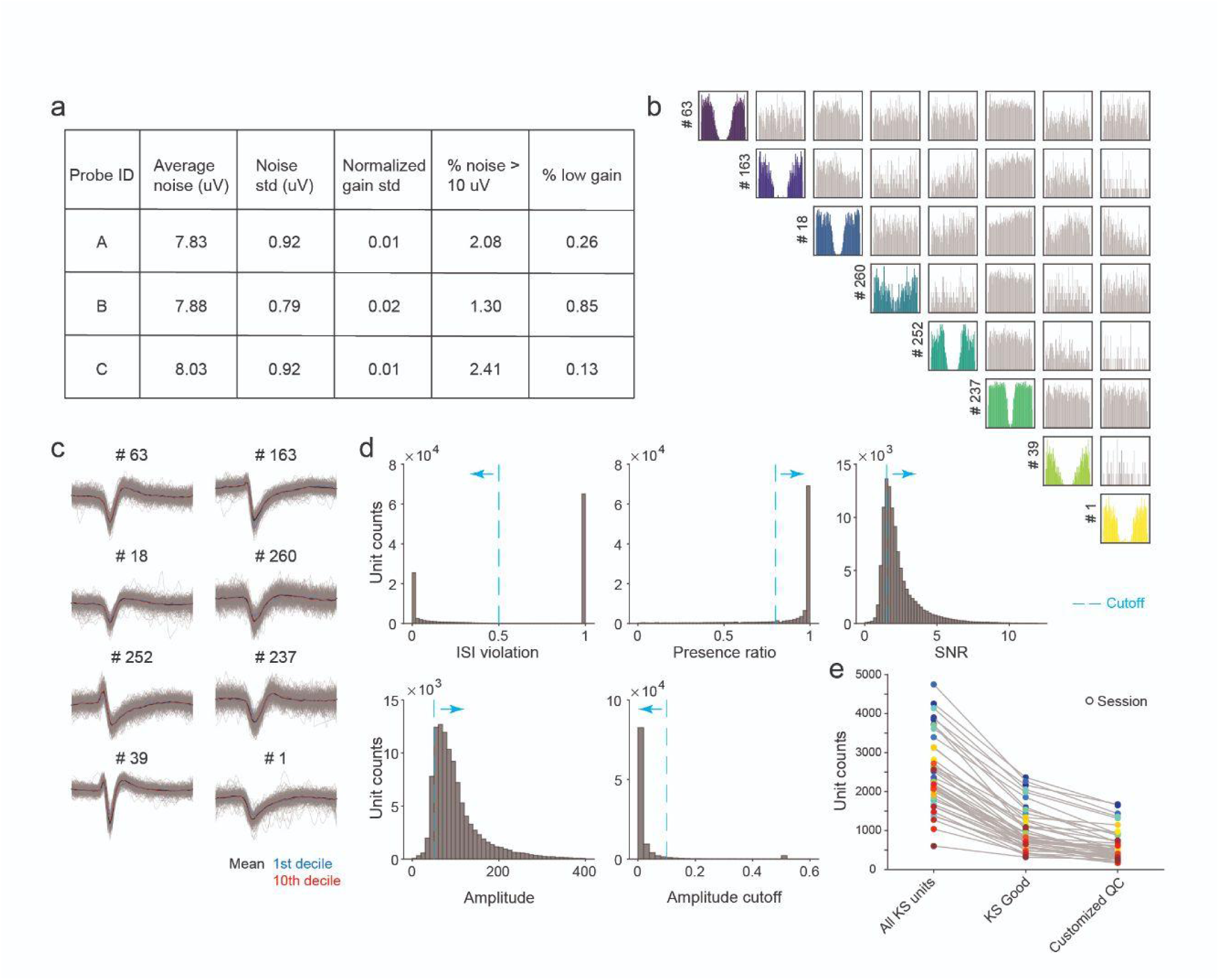
Probe characterization, unit quality metrics, and selection criteria. **a,** Statistical summary of noise and gain metrics in saline. “Low-gain” channels are defined as those with a gain value below 50% of the average; these channels were excluded from all other measurements. **b**, Representative auto- and cross-correlograms of units in a recording session. Examples of autocorrelograms (ACGs) for isolated units and cross-correlograms (CCGs) between unit pairs following spike sorting. The clear refractory periods in the ACGs indicate high-quality single-unit isolation with minimal multi-unit contamination. **c**, Stability of unit waveforms. Waveform profiles for the example units shown in (**b**). Each panel displays 300 randomly sampled spike waveforms (light grey) superimposed with the session mean (black). To demonstrate temporal stability, mean waveforms from the first decile (blue) and last decile (red) of the recording session are overlaid. The high degree of overlap between the early and late waveforms confirms the stability of the extracellular recordings and the consistency of unit isolation throughout the session. **d**, Distribution of single-unit quality metrics. Histograms representing the distribution of metrics for all units identified by Kilosort. Cyan dashed lines indicate the customized thresholds applied to isolate high-quality single units. The criteria are: (1) inter-spike-interval (ISI) violations within a 1.5-ms window must be < 0.5 false positive rate; (2) spike presence must exceed 80% of the total session duration; (3) the signal-to-noise ratio (SNR) must be > 1.5; (4) mean unit amplitude must exceed 50 µV; and (5) the amplitude cutoff must be < 0.1 fraction of missing spikes. **e**, Comparison of unit yields across quality control categories. The categories include: (1) all raw units identified by Kilosort; (2) units automatically labeled as “good” by the Kilosort algorithm; and (3) units meeting our customized quality control criteria (as defined in **d**). The use of the NP QB platform significantly maintains high unit yields even after the application of these stringent customized filters.

**Figure S2:**
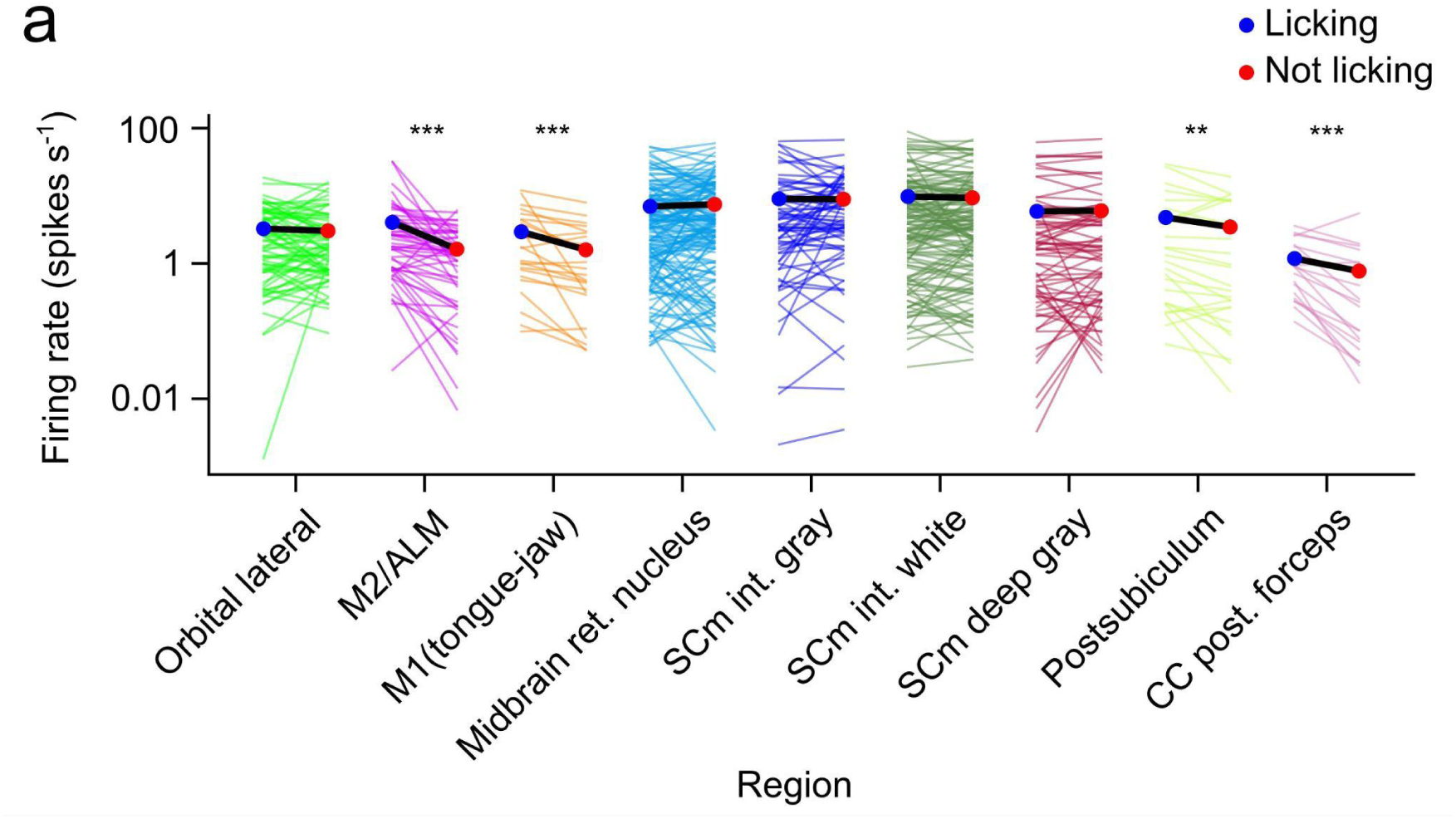
Firing rates of units shown in Figure 4 during Licking and Not licking periods. **a,** Mean firing rates of units from different regions during the Licking and Not licking periods. Black lines show region-wise means. Two-sided Wilcoxon signed-rank test, *** p < 0.001, ** p < 0.01

**Figure S3:**
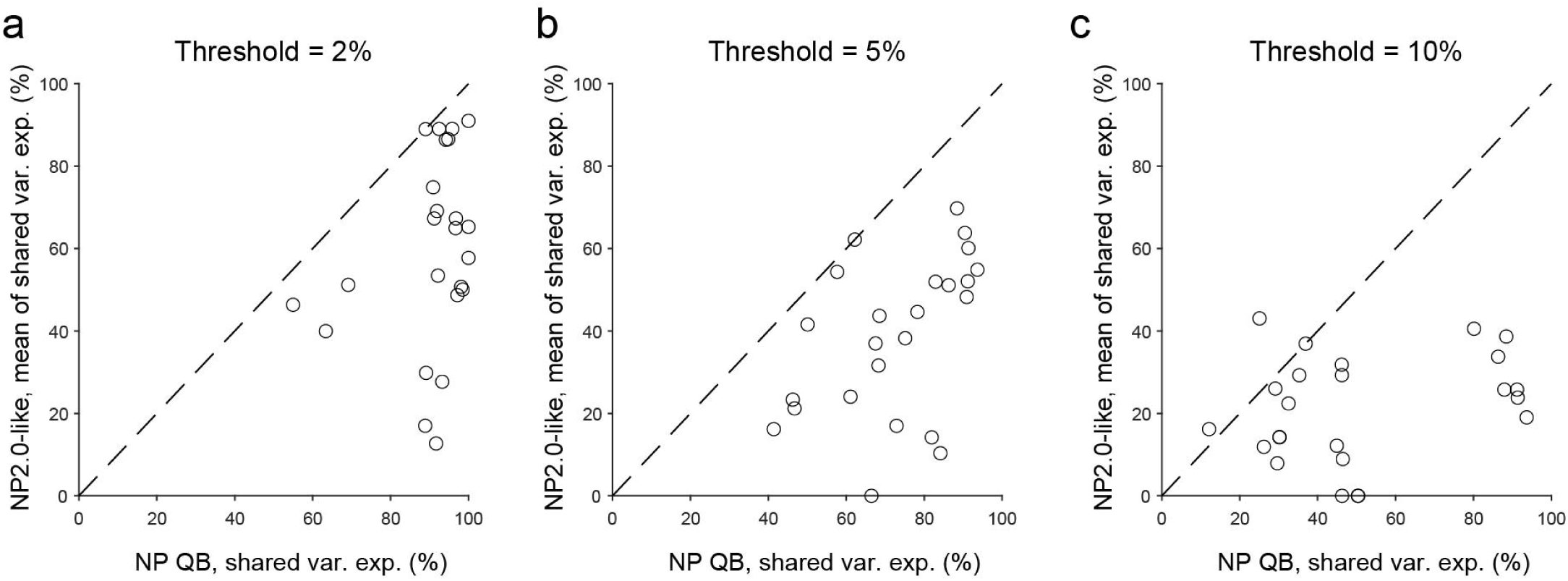
Shared variance explained of NP QB and NP2.0-like probes at different variance thresholds. Shared variance explained from mDLAG analysis for NP QB probes is plotted against the mean of variance explained across the corresponding NP2.0-like simulations. Each plot shows analysis for a different variance threshold (2% in **a**, 5% in **b** and 10% in **c**).

**Figure S4:**
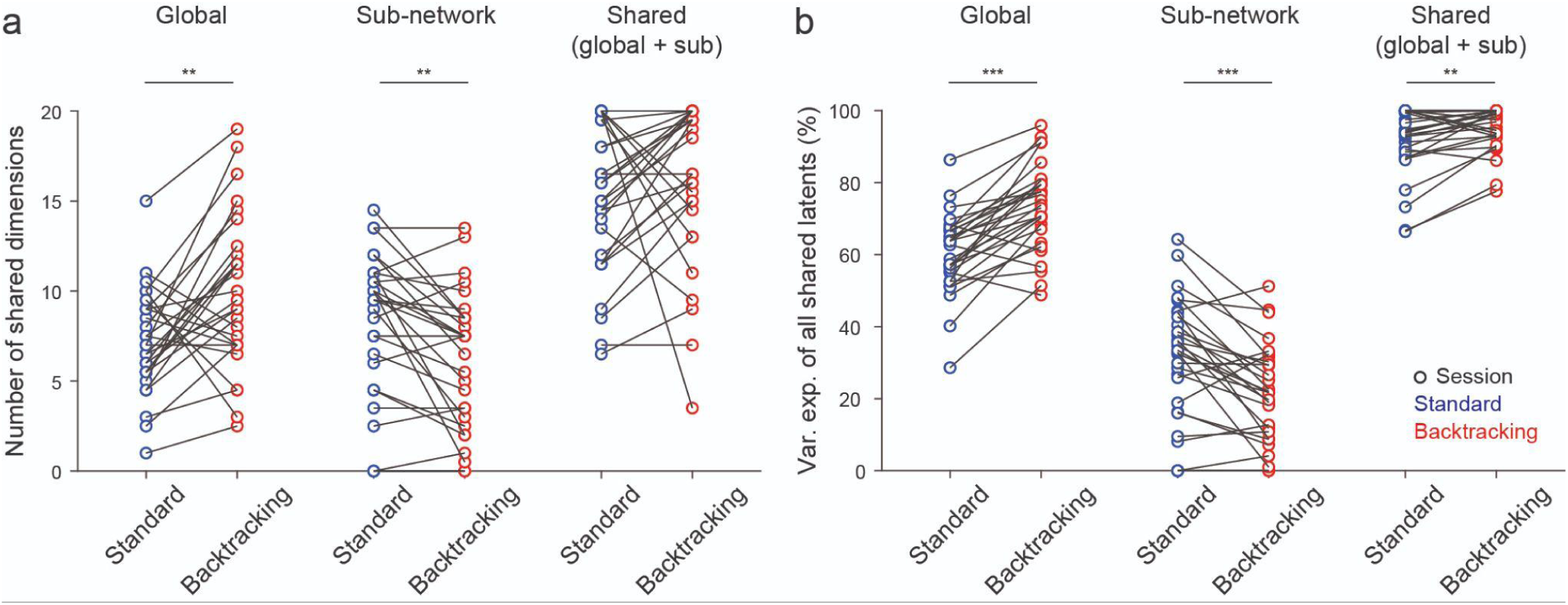
Comparison of latent dimensions and explained variance for standard and backtracking trials. **a**, Number of latent dimensions identified with mDLAG analysis during standard (blue) and backtracking (red) trials. While the total number of shared dimensions (comprising both global and sub-network components) remains consistent across trial types, backtracking trials exhibit a significant shift in latent structure: global dimensions increase in number, while sub-network dimensions decrease. **b**, Explained variance of latents in standard and backtracking trials. Total variance accounted for by all shared latent dimensions is significantly higher during backtracking trials compared to standard trials. This increased explanatory power is driven primarily by the global latent component.

**Table S1:**
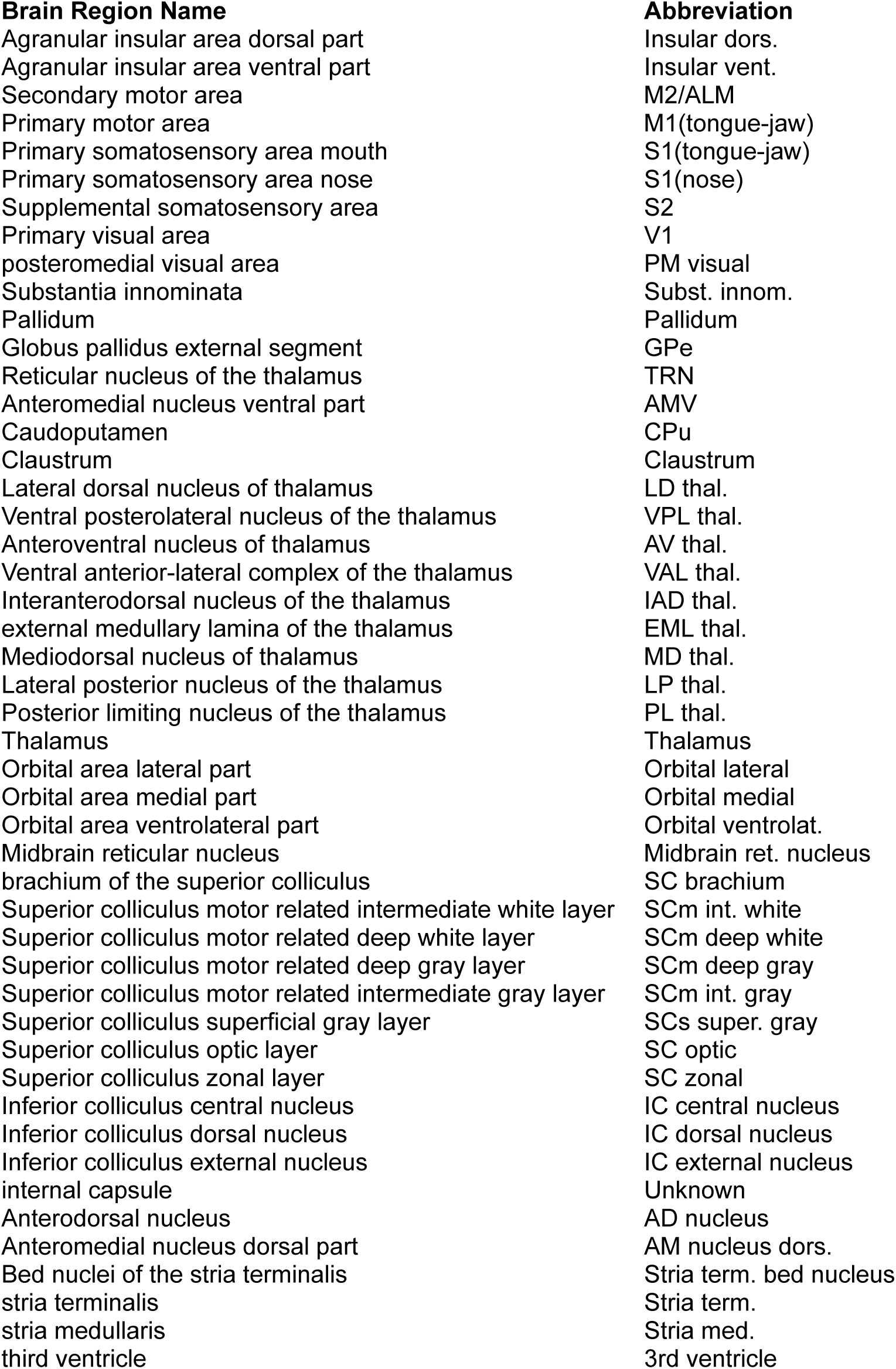
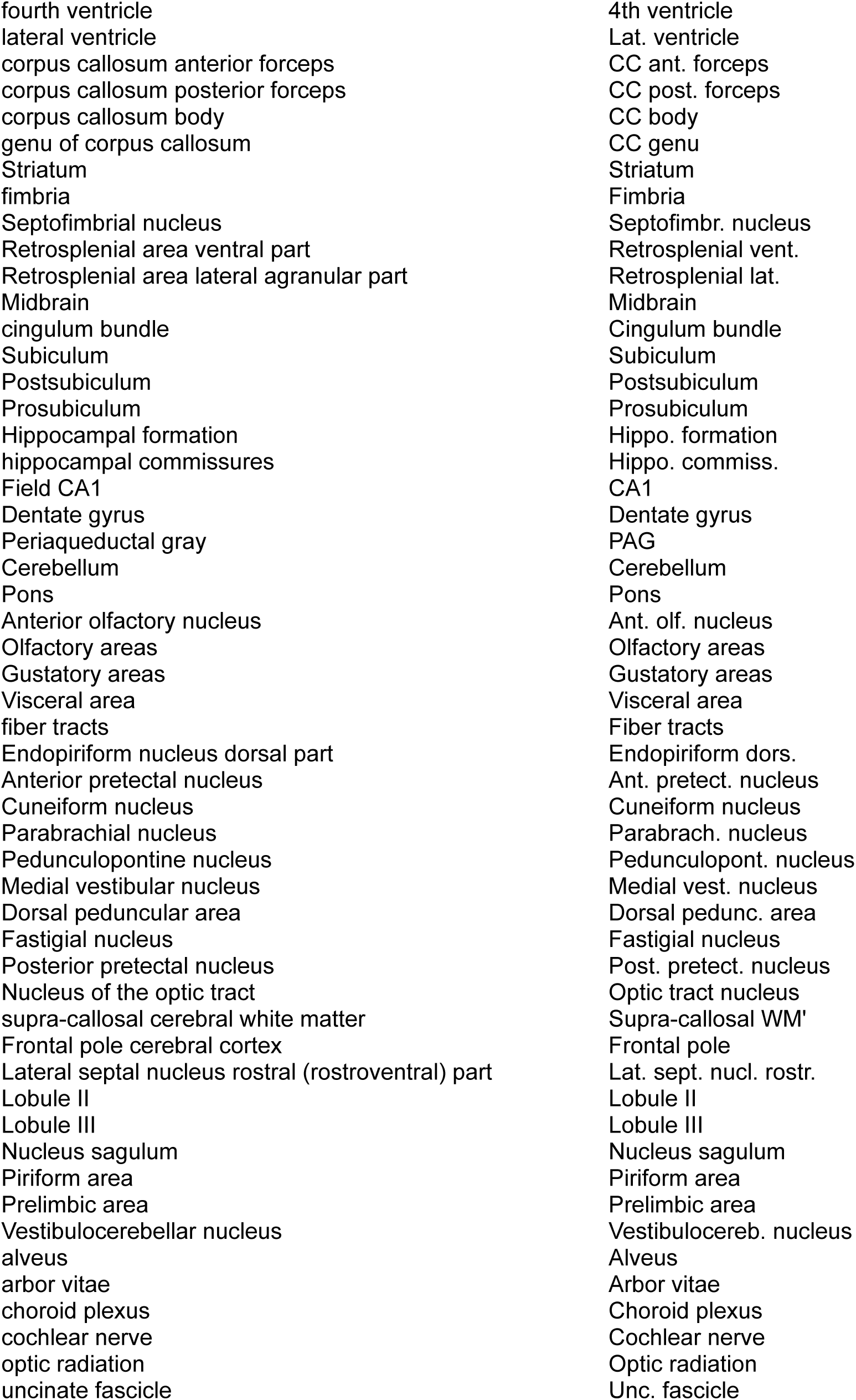
Full brain region names and corresponding abbreviations. Probe tracks were reconstructed using SHARP-Track, and histological slice images were registered to the Allen Mouse Brain Common Coordinate Framework (CCF). This table maps the brain region names from the CCF to the corresponding abbreviations used in this study.

## Star Methods

### Mice

All experimental procedures were performed in accordance with protocols approved by the Johns Hopkins University Animal Care and Use Committee. Six wild-type mice (3 male, 3 female), aged 2–3 months at the start of training, were included in the study. Animals were housed in a vivarium maintained on a 12-hour reverse light-dark cycle. Mice were housed in groups prior to surgery and transitioned to single housing following surgical procedures and throughout the duration of behavioral experiments.

### Surgery

To enable head-fixed behavioral training and subsequent electrophysiological recordings, mice were prepared with a headpost implantation and a clear-skull cap (Citation).^30^ To accommodate various brain targets, the headpost was positioned posteriorly over the cerebellum in four mice and more anteriorly over Bregma in two mice. Surgical anesthesia was maintained with isoflurane (1-2% in O2; Surgivet) while the mice were secured in a stereotaxic frame (David Kopf Instruments). To maintain stable body temperature throughout the procedure, animals were placed on a homeothermic heating blanket (Harvard Apparatus). Local analgesia was provided via subcutaneous lidocaine (2%, VetOne), and systemic inflammation was managed with an intraperitoneal injection of ketoprofen (Covetrus, diluted to 1 mg/ml). Following the removal of the scalp and periosteum, headposts were secured using a dental adhesive system (C&B Metabond; Parkell). The remaining exposed skull was made transparent by applying a thin layer of clear dental cement followed by a coating of cyanoacrylate glue (Krazy Glue). To maintain the integrity of the clear-skull preparation and prevent scratching, the surface was protected with a silicone elastomer (Kwik-Cast) between experimental sessions.

To guide dual-probe insertions, recording trajectories were planned using Pinpoint.^31^ Prior to electrophysiological recording, small craniotomies (approximately 1 x 1.5 mm) were performed at the coordinates specified in the electrophysiology section. The overlying dental acrylic and skull were carefully thinned using a dental drill, and the remaining bone was removed using a fine tungsten needle or surgical forceps to expose the dura. As recordings were completed in initial targets, additional craniotomies were performed in novel locations to access further brain regions. Across the six mice included in this study, a total of 6–9 craniotomies were performed per animal to facilitate comprehensive mapping.

### Sequence licking task

#### Task control

Mice performed sequential, directed licking toward a moving lick port, as previously described in Xu et al.^15^ A stainless-steel lick port was mounted on two-axis linear motors (LSM050B-T4 and LSM025B-T4, Zaber Technologies), and port height was adjusted using a manual linear stage (MT1/M, Thorlabs). Task control was implemented on a Teensy 3.1 microcontroller using custom MATLAB (R2020b) and Arduino (1.8.13) software. Lick contacts were detected using a custom lick-detection circuit (Janelia Research Campus). During sequence execution, lick contacts triggered port movements between predefined locations along an arc. At the end of each trial, a water reward was delivered via a solenoid valve (LHDA0531415H, The Lee Company).

#### Behavioral training

All behavioral experiments were performed with head-fixed mice during the dark phase. One week prior to training, mice were water restricted and received 1 mL water per day until body weight stabilized at ∼80% of baseline. On training days, mice performed the task until satiety (typically ∼1 h/day). Body weight was measured before and after each session to estimate water consumption, and supplemental water was provided if intake during training was <0.6 mL to maintain stable body weight. On non-training days, mice received 1 mL water.

Mice were first trained to perform standard sequences. Each trial began with an auditory go cue (pure tone; 15 kHz for RD079 and 8 kHz for all other animals), after which mice executed a sequence of licks toward the lick port and received a water reward at the end of the sequence. Seven lick-port positions were defined along an arc centered on the estimated root of the tongue, with arc radius corresponding to tongue-to-port distance. Training began with a smaller radius (e.g., ∼4 mm) and narrower angular spacing (e.g., 60°, 70°, 80°, 90°, 100°, 110°, 120° for L3, L2, L1, Mid, R1, R2, R3). As mice improved in modulating tongue direction, the radius and/or angular spacing were gradually increased (e.g., radius to ∼4.5 mm; angles to 45°, 60°, 75°, 90°, 115°, 130°, 145° for L3–R3), such that the first and final positions spanned a wide range about the midline. The next trial went through the same port locations but in the opposite direction.

Mice were considered proficient at standard sequences once they could reliably target each port position with minimal missed licks across trials. After standard proficiency, backtracking sequences were introduced, where the lick port moved backwards after the lick at Mid position. Behavioral and/or neural recordings began after mice had reached a criterion level of performance on backtracking sequences (e.g., at least one-third of backtracking trials completed with only 1–2 missed licks following the Mid touch).

### Quantification of behavioral variables

#### High-speed videography

Mouse behavior was recorded using high-speed videography at 400 frames/s with hardware-triggered acquisition of both bottom and side views of the face. The face was illuminated with an 850-nm LED (LED850-66-60, Roithner LaserTechnik). Videos were acquired through a 0.25× telecentric lens (Edmund Optics) using a PhotonFocus DR1-D1312-200-G2-8 camera and Streampix 7 software (Norpix). Videos were saved separately for each trial with a maximum duration of 20 seconds; trials exceeding the maximum duration were excluded from analysis.

Tongue landmarks were tracked using DeepLabCut^32^ by fine-tuning ImageNet-pretrained ResNet-50 convolutional neural networks to annotate the tongue base, tongue tip, and jaw tip in each frame from both bottom and side views.

#### Tongue kinematics and task parameters

Tongue azimuthal angle (θ) was defined in the bottom view as the angle between the vertical axis and the line connecting the tongue base and tongue tip; by convention, negative values indicate leftward licking and positive values indicate rightward licking. Tongue length (L) was defined as the Euclidean distance between tongue base and tongue tip in the bottom view. Frames in which tracked landmarks were not visible (e.g., when the tongue was inside the mouth) were excluded using the DeepLabCut confidence score and additional geometric constraints. Specifically, tongue-out frames were defined as those with confidence exceeding a threshold (typically 0.8) and, when applied, with tongue length within a plausible range (e.g., 0.9–4.8 mm). To reduce frame-to-frame jitter in landmark estimates, kinematic traces were smoothed using median and Savitzky–Golay filters.

Additional task variables included tongue speed (L′), relative sequence time (τ), and lick phase. Tongue speed was computed as the time derivative of tongue length. Relative sequence time was defined within each trial as a normalized time variable ranging from 0 at go cue to 1 at the last drive lick (which triggered the water reward). Lick phase was derived from the jaw-tip z coordinate, with 0–π corresponding to the opening phase and π–2π corresponding to the closing phase.

### Electrophysiology

Neuropixels Quad Base probes were coated with either CM-DiI (Invitrogen by Thermo Fisher Scientific, V22888) or DiD (5-10 mg/mL, Invitrogen by Thermo Fisher Scientific, D307) to facilitate post-hoc histological verification of recording sites. Dual-probe configurations were employed to target multiple sensory and motor regions simultaneously. For S1 (tongue-jaw), probes were inserted at 0.5 mm anterior and 3.8 mm lateral to bregma, reaching a depth of 1.3 mm at a 40° angle. Additional targets included M1 (tongue-jaw) (1.5 mm anterior, 2.5 mm lateral, 1.3 mm depth, vertical), ALM (2.5 mm anterior, 1.5 mm lateral, 1.3 mm depth, vertical), caudate putamen (CPu) (0.5 mm anterior, 3.8 mm lateral, 3.75 mm depth, vertical), and superior colliculus (SC) (3.5 mm posterior, 1.5 mm lateral, 2.5 mm depth, vertical). To achieve simultaneous recording of the SC and midbrain trigeminal nucleus (MEV), the insertion point was shifted to 3 mm posterior and 1.5 mm lateral, reaching 4 mm depth at a 21° angle on the sagittal plane. For concurrent motor area and CP recordings, the probes reached the CP from the contralateral side (0 mm anterior, 1 mm lateral, 6.4 mm depth, 39° angle). Following insertion, the brain was covered with a layer of 1.5% agarose (Sigma-Aldrich), with an approximately 10-minute recovery period allowed prior to data acquisition.

### Histology and probe tracking

After the completion of experiments, mice were transcardially perfused with 4% paraformaldehyde in phosphate buffered saline. The brains were extracted and fixed overnight. 100 μM slices were obtained using an EasySlicer (Pelco easiSlicer, Ted Pella) and imaged (BX-41, Olympus).

Slice images were registered to the Allen Mouse Brain Common Coordinate Framework (CCF) and probe tracks reconstructed using SHARP-Track (github.com/cdoi.org/10.1101/447995ortex-lab/allenCCF).^33^

### Data preprocessing

Neural signals and behavioral timestamps were acquired using Neuropixels OneBox (ONEBOX_1000) and PXIe (PXIE_1000) systems. Each probe was connected to its respective acquisition system via independent headstages and cables, with SpikeGLX software used to configure recording settings and channel selection. Post-acquisition, data were processed through a unified electrophysiology pipeline for spike sorting via Kilosort4^14^. Units were retained for decoding (Figure 3), Granger causality (Figure 4), and mDLAG (Figure 5) analyses based on quality metrics adapted from the Allen Institute: inter-spike-interval (ISI) violations within a 1.5 ms window were required to be < 0.5 false positive rate, spike presence was required for > 80% of the session, unit amplitude exceeded 50 µV, amplitude cutoff was < 0.1 fraction of missing spikes, and the signal-to-noise ratio (SNR) was > 1.5.

For downstream analysis, neural spike rates were binned at 25 ms for mDLAG whole-sequence analysis (Figures 5d–g) and at 5 ms for mDLAG back-tracking analysis (Figures 5h–j). All spike rates were smoothed with a 15 ms Gaussian kernel. Finally, to improve trial-to-trial consistency during mDLAG analysis, data were temporally morphed according to the framework established by Xu et al.^15^

### NP2.0-like dataset generation

To generate an “NP2.0-like” control dataset, we computationally resampled the NP QB data (1,536 channels) to mimic the specific channel map and recording capacity of a standard NP2.0 probe (384 channels; Figures 1, 4 and 5). Eight distinct channel configurations were simulated to evaluate the impact of spatial sampling. Four simulations were dedicated to single-shank configurations (Shanks 0, 1, 2, and 3), each utilizing 384 channels per shank. The remaining four simulations modeled multi-shank “banks” across all four shanks, spanning the probe from bottom to top in vertical tiers: Bank 0 (channels 0–96), Bank 1 (channels 97–192), Bank 2 (channels 193–288), and Bank 3 (channels 289–384). This standardized resampling approach allowed for a direct comparison between NP QB and NP2.0-like conditions.

### Task variable decoding

To validate the quality of recordings, we linearly decoded task and behavioral variables from the population neural activity in various brain regions using Ridge regression, separately for each session. In each trial, we considered the one second period centered on the time of lick port contact at the Mid position and extracted all frames in that period where the tongue was outside the mouth. To ensure fair comparisons of decoding performance between variables, we ignored tongue-in-mouth frames for all variables, including ones that could be tracked continuously. For each variable, values from all relevant frames were concatenated across trials into a vector *v* ϵ ℝ^*T*^ (*T* total frames). The optimization problem solved by Ridge regression was then

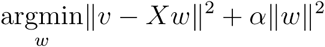

where *X* ϵ ℝ^*T×N*^ (*N* total units from this region and session) was the neural firing rate data matrix, *ω* was the extracted latent vector of weights, and α controlled the strength of regularization. The neural data matrix was then projected on the latent weights to obtain predictions, and decoding performance was quantified as the coefficient of determination of the predicted variable. We used 10-fold cross-validation to choose the optimal value for *α* for each session from among seven candidate values. In all places, we report decoding performance for a completely held-out dataset (excluded from the cross-validation set used to arrive at *α*).

### Trial classification

We used linear support vector machines (linear-SVM) to quantify the discriminability between population activity vectors in standard and backtracking trials. We trained separate classifiers for each time bin in each session, using vectors *r* ϵ ℝ^*N*^ of firing rates of units recorded from a given region as the data-points.

We regularized linear-SVM models to reduce overfitting. The regularization parameter C (inversely proportional to the strength of regularization) was chosen from among seven candidate values (same as for decoding models above) using 10-fold cross-validated classification performance. Since the numbers of standard and backtracking trials were unequal in every session, C was proportionally weighted to be larger for the backtracking trials. In all places, we report classification performance (proportion of correctly classified trials) for a completely held out set of trials (excluded from the cross-validation set used to arrive at C).

### Granger causality

Granger causality measures whether past values of an independent variable have any predictive power over future values of a dependent variable. More specifically, it asks whether we can significantly improve the predictions of future values of the dependent variable by taking into account the independent variable, compared to the case where we consider past values of the dependent variable alone.

For each pair of neurons under consideration, we calculated Granger causation in both directions, i.e., with each neuron in the pair serving as the independent or dependent variable. Firing rates of the dependent unit lagged the independent one with a delay of a single time-bin (2.5 ms). A Granger causal link was deemed to exist only if significant Granger causation was found in both directions (chi-square test with Bonferroni correction for multiple comparisons for a family-wise *α* of 0.05).

### mDLAG modeling and latent analysis

To identify shared signals across multiple brain regions, we applied multigroup Delayed Latent Across Groups (mDLAG) modes.^19,34^ We used a frequency-domain acceleration approach to fit the models efficiently.^20^

To ensure sufficient statistical power, brain areas were only included in the analysis if they contained at least 20 recorded single units. For each model, we specified 20 shared latent dimensions. These shared latents were ranked according to the mean variance explained across all included brain areas. A variance-explained cutoff was applied to determine the significance of a latent’s contribution to a specific brain area. Shared latents were categorized into two types, global and sub-network latents. Global latents contribute significantly to the variance in all included brain areas, while sub-network latents contribute significantly to at least two areas, but fewer than the total number of areas in the model.

To identify shared latents during the sequence licking task, we analyzed a time window from -0.65 to +0.85 seconds relative to the middle lick, capturing the full sequence (Fig. 5d). Neural data were temporally morphed to ensure consistency across trials. Right-to-left and left-to-right sequences were modeled independently. To investigate neural network dynamics during motor program reorganization, we compared standard versus backtracking trials. This analysis focused on the 0.1 to 0.4-second window following the middle lick, where motor plan updates primarily occur (Fig. 5h). For the NP2.0-like simulations, neurons were sub-sampled according to specific NP2.0 channel maps (see Methods). To ensure a direct comparison, all mDLAG hyperparameters were kept identical between the NP QB and NP2.0-like conditions.

## Acknowledgements

This work was supported in part by NIH BRAIN Initiative Grant U01NS115587 and 1U19NS137920-01.

